# Temporal Dynamics of Parvalbumin Interneurons and Perineuronal Nets During Learned Helplessness

**DOI:** 10.64898/2026.09.16.752101

**Authors:** Labiba Aziz, Suli Wang, Michele Assef, Dani Dumitriu, Alessia Manganaro

**Affiliations:** Department of Pharmacological Sciences, Renaissance School of Medicine, Stony Brook University, Stony Brook; Center for Early Relational Health, Columbia University, New York, NY, United States; Division for Child and Adolescent Health, Department of Pediatrics, Columbia University Irving Medical Center, New York, NY, United States; Division of Developmental Neuroscience, Department of Psychiatry, Columbia University Irving Medical Center, New York, NY, United States

## Abstract

This study followed the recruitment of medial prefrontal cortex parvalbumin (PV^+^) interneurons and their perineuronal nets (PNN) across the two stages of the learned helplessness (LH) paradigm: the initial uncontrollable stress experience and the later expression of helpless or resilient coping. Combining TRAP2-based activity mapping with a staggered-endpoint design, three main findings emerged on the functional and structural neuronal correlates in prelimbic (PL) and infralimbic (ILA) subregions during LH paradigm. First, prefrontal recruitment during the first inescapable shock session already predicted the coping strategy that would be expressed two days later during escape avoidance test, before any behavioral divergence was observable. Second, the expression of parvalbumin protein (PV) in this cortical subpopulation of interneurons followed a dynamic trajectory rather than a static stress change, transiently reduced across all shocked animals at the escape test, and back to baseline over the following week in both PL and ILA, except in animals that went on to be classified as “helpless”. Third, PNN intensity temporal patterns did not track stress exposure in either subregion, yet PNN and PV became correlated with one another specifically in ILA in stressed animals, pointing to a stress-induced structural coupling rather than a constitutive relationship between the two markers.

## Introduction

Stress-related psychiatric disorders such as major depression and post-traumatic stress disorder are among the leading causes of disability worldwide, yet only a subset of individuals exposed to severe or uncontrollable stress go on to develop them, while others remain resilient^1,2^. Identifying the neural mechanisms that separate vulnerability from resilience is therefore central to understanding who is at risk, and to defining targets for intervention rather than treating symptoms once they are established. Preclinical models that segregate vulnerable from resilient individuals following an identical stressor provide a tractable way to isolate these mechanisms ^3–11^. Learned helplessness (LH) separates the experience of uncontrollable stress from the later behavioral response to a controllable stressor ^12–14^. In the classic rodent paradigm, animals first receive inescapable shock (IS) and are subsequently tested with escapable shock (ES). Although all animals initially experience the same uncontrollable stressor only a subset later fail to initiate escape responses and are classified as “helpless,” while animals that actively avoid the stressor are classified as “resilient.” LH can therefore be used to examine two distinct stages: the emergence of vulnerability during uncontrollable stress and the later expression of helpless or resilient behavior ^14,15^.

Early LH studies emphasized hippocampal mechanisms involved in encoding the uncontrollable stress experience ^16–19^, including stress-associated remodeling of local inhibitory circuits ^20–23^. The later decision to initiate or suppress an escape response, by contrast, is governed by prefrontal circuits: rat studies have established a pivotal role for the medial prefrontal cortex (mPFC) in the direct control of goal-directed behavior and action selection and have further implicated the mPFC in both the detection of behavioral control and the expression of helplessness ^24–26^.

In mice, prefrontal PV⁺ interneurons have been linked more directly to this process: excitatory synaptic drive onto mPFC PV⁺ interneurons is weakened specifically in helpless animals, and pharmacogenetic suppression of these interneurons during the paradigm promotes helpless behavior, indicating that reduced PV⁺ interneuron engagement contributes to helplessness^27^. Despite this, it remains unclear from the existing literature at what stage of the paradigm helplessness emerges — whether prefrontal inhibitory circuit differences arise during the initial uncontrollable experience or only become apparent once helpless behavior is expressed — an ambiguity that single-timepoint studies cannot resolve. We hypothesized that this individual variability emerges early, during the initial inescapable stress exposure itself, and is already reflected in prefrontal recruitment and inhibitory circuit dynamics before any behavioral divergence is observable, rather than arising only when helpless or resilient behavior is later expressed.

We focused on the prelimbic (PL) and infralimbic (ILA) regions of the mPFC, also referred to as dorsal and ventral mPFC respectively, which are both involved in behavioral responses to stress^28–30^. Within these regions, PV⁺ interneurons provide strong inhibition of pyramidal neurons and help regulate the timing and coordination of prefrontal activity ^31–34^. By controlling which pyramidal-cell ensembles are recruited, PV⁺ interneurons may shape how the mPFC responds to uncontrollable stress and later escape opportunities. Consistent with this idea, manipulating prefrontal PV⁺ activity can alter stress-related behavior ^27,35^.

PNNs are extracellular matrix structures that preferentially surround PV⁺ interneurons and regulate their stability and capacity for plasticity. Although PNNs stabilize inhibitory circuits, they remain dynamic in the adult brain, where reduced PNN expression permits greater plasticity. Conversely, chronic stress can increase perineuronal net and extracellular matrix deposition around PV⁺ interneurons, and this accumulation constrains plasticity and impairs memory, an effect reversed by enzymatic degradation of the excess matrix ^43,44^. PV⁺ interneuron and PNN abnormalities are reported across stress-related and other psychiatric conditions, including depression, PTSD, and schizophrenia, and restoring normal PV⁺/PNN function has been proposed as a therapeutic avenue^34,37,39^. Examining PV⁺ interneurons and PNNs across LH may therefore reveal how prefrontal inhibitory circuits change between the emergence and expression of helplessness.

To determine when these differences emerge, we used TRAP2 (Targeted Recombination in Active Populations)^40^ system to permanently label neuronal ensembles recruited during either the initial uncontrollable stress experience or subsequent escape testing. Unlike immediate-early-gene staining, which captures only the activity present at the moment of tissue collection, TRAP2 provides a permanent, activity-dependent tag of the neurons active within a defined behavioral window, allowing us to relate the ensemble recruited during uncontrollable stress to a coping phenotype that emerges only days later. Rather than sampling a single endpoint, we then tracked PV⁺ interneurons and PNNs across successive stages of the paradigm, from the initial inescapable shock through the escape test and into the following week. This temporal approach was designed to distinguish differences that are present from the outset from those that emerge, resolve, or persist over time, and thereby determine whether helplessness and resilience reflect differences that arise during uncontrollable stress, during behavioral expression, or through dynamic remodeling between these stages.

## Materials and Methods

This study was designed to characterize inhibitory circuit dynamics across distinct stages of learned helplessness (LH) by permanently labeling neuronal ensembles recruited during inescapable shock (IS) or escapable shock (ES) and validating these labeling timepoints using independent perfusion cohorts.

A total of 59 TRAP2:Ai14 mice were used to permanently label neurons activated during specific phases of the LH paradigm:

1. IS1 group (n =24): Mice received one intraperitoneal injection of 4-OHT soon after IS ats day 1 to label neurons activated during the initial uncontrollable stress exposure.
2. ES group (n =35): Mice received one intraperitoneal injection of 4-OHT soon after ES testing to label neurons recruited during the escape behavior testing, when helpless and resilient phenotypes are expressed.

For the matched behavioral–histological analyses, animals were excluded based on quality criteria for brain slice registration and tissue and staining quality (high background or missing one of the marker investigated), resulting in a final sample of 17 mice per group: for IS1 group 6 controls (4 females and 2 males) and 11 shocked mice (1 female and 10 males), for ES group 6 controls (4 females and 2 males) and 11 shocked mice (5 females and 6 males).

To validate TRAP labeling windows and assess perfusion timing independently of tdTomato stabilization kinetics, an additional 54 wildtype C57BL/6J mice were used. Of these, 60 mice were perfused at specific and distinct timepoints corresponding to first unescapable stress exposure, escape testing, or post-stress recovery.

1. Home cage controls (HC; n = 6): Mice were perfused without prior exposure to LH or ASDS to establish baseline measures.
2. LH Day 1 group (n = 16): Mice were perfused one hour after LH Day 1 training to validate neuronal activation captured in the LHD1 TRAP2:Ai14 cohort (3 control females, 3 control males, 5 shocked females, 5 shocked males).
3. LH Day 3 group (n = 16): Mice were perfused one hour after LH Day 3 testing to validate neuronal activation captured in the LHD3 TRAP2:Ai14 cohort (3 control females, 3 control males, 5 shocked females, 5 shocked males).
4. LH +7 days group (n = 16): Mice were perfused ten days after LH Day 3 testing to control for tdTomato stabilization delay in TRAP2:Ai14 mice and to assess longer-term effects of LH exposure.

### Animals

10-12 week-old male and female mice were used in this study. Wildtype C57BL/6J mice were obtained from Jackson Laboratory (strain #000664) and received 3 weeks before start of experiment to allow habituation. The double transgenic TRAP2:Ai14 mice were bred in-house by crossing the homozygous TRAP2 line Targeted Recombination in Active Populations (Fostm2.1(icre/ERT2)Luo/J, Jackson Laboratory #030323) with the homozygous Cre-reporter line Ai14 (B6.Cg-Gt(ROSA)26Sortm14 (CAG-tdTomato)Hze/J, Jackson Laboratory #007914).

Both wildtype and TRAP2:Ai14 mice were group-housed 3-5 per cage same sex under a 12-hour light/dark cycle (lights on 7 am–7pm) with ad libitum food and water in a temperature-controlled room (18–23°C) with 40–60% humidity a week before the acclimation for the experiment started. Randomly assigned experimental and control animals were all housed adjacently in the same room. Cage-mates were not changed throughout the experiment. Experimental procedures and testing were performed during the light cycle between 12-3 pm in procedural rooms close to the housing. All experiments were conducted under the regulations of Columbia University and the New York State Psychiatric Institute Institutional Animal Care and Use Committee (IACUC).

### Learned Helplessness Procedure

During the acclimation period, mice were handled for two days, followed by five consecutive days of intraperitoneal saline injections, administered at the same time of day as the upcoming LH procedures. Both experimental and control groups underwent this protocol to control for injection-related stress. As shown in Figure 1, LH was performed using LH equipment (an automated shuttlebox with two adjacent compartments, Med Associates) and a PC that executed a custom-made coded script and collected experimental readouts via software M5 (Med Associates Inc). LH boxes deliver an electric shock via metal grid floors and collect movement data via infrared sensors. The isolated boxes also have an automated gate that allows mice to move between the two compartments when opened.

**Figure 1:**
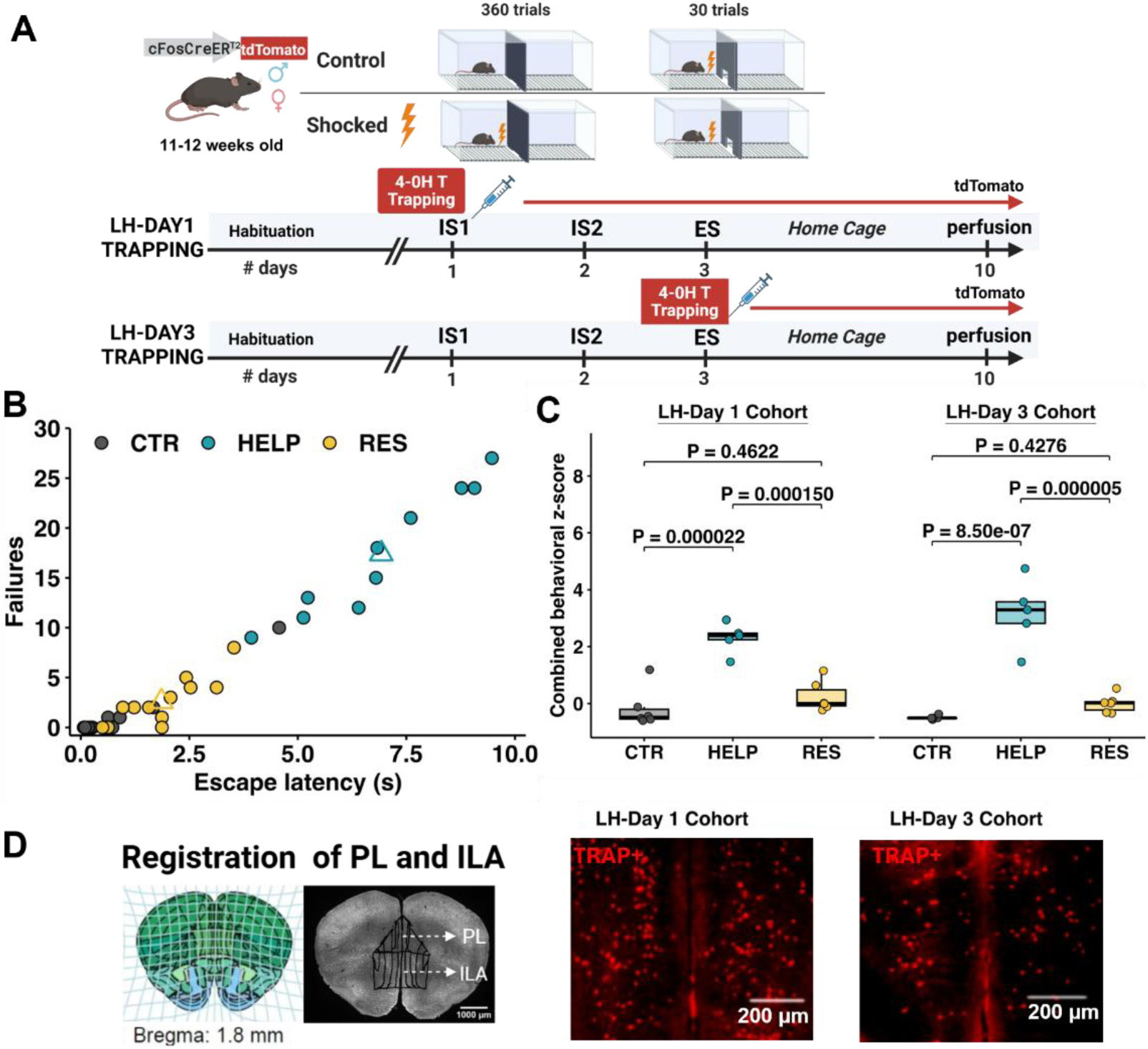
Behavioral paradigm and behavioral classification in the LH paradigm. **(A)** Experimental timeline for activity-dependent labeling in TRAP2 mice. Control mice underwent equivalent shuttle-box exposure without inescapable shock, whereas shocked mice received inescapable shock on days 1 and 2. All mice completed the 30-trial escape test on day 3. Neurons active during the first inescapable-shock session (IS1; LH-Day 1 cohort) or the escape session (ES; LH-Day 3 cohort) were permanently labeled by administration of 4-OHT, and tissue was collected on day 10. **(B)** Behavioral classification based on escape failures and mean escape latency, k-means clustering (k = 2) was performed on the pooled stress-exposed animals from both cohorts. Non-shocked controls were displayed but excluded from clustering. Circles represent individual animals classified as control (CTR; gray), helpless (HELP; teal), or resilient (RES; yellow). Open triangles indicate the constrained cluster centroids: HELP, 17.40 failures and 6.92 s latency; RES, 2.58 failures and 1.86 s latency. CTR, n = 12; HELP, n = 10; RES, n = 12. **(C)** Combined behavioral z-scores derived from standardized escape failures and escape latency are shown separately for the LH Day 1 and LH Day 3 cohorts. Higher scores indicate more severe helplessness-like behavior. Points represent individual animals; boxes indicate the median and interquartile range, and whiskers extend to 1.5 times the interquartile range. **(D)** Registration of the prelimbic (PL) and infralimbic (ILA) regions at approximately 1.8 mm anterior to bregma and representative tdTomato fluorescence images from the two cohorts. The dashed line indicates the boundary between PL and ILA. Scale bars = 200 pm.

LH was executed similarly to previously described protocols ^14,41^. On Days 1 and 2 (IS1 and IS2 training), mice in the experimental group (LH-shocked or STR) received 360 inescapable foot shocks (0.3 mA, randomized 1–3 s duration, randomized 1–15 s ITIs) corresponding to approximately 55-minute session. Control mice were placed in the boxes for the same amount of time but did not receive shocks. Each session began with 250-second habituation. On Day 3 (testing), all mice received escapable foot shocks. First, mice were allowed to habituate by moving through both chambers for 120 seconds before the door closed and trials started. Each trial began with gate opening and a 0.3 mA shock. If the mouse moved to the adjacent compartment within 10 seconds, the shock session ended, the door closed and the escape latency was recorded by the program. Failure to escape the shock within 10 seconds was considered a failed shock trial and a 10 sec latency will be recorded. Each animal had 30 test trials on Day 3. LH boxes were thoroughly cleaned with odor-neutralizing and sterilizing wipes between animals. The mice were transferred back to their home cage with their housemates after each testing day.

Two separate groups of TRAP2:Ai14 mice received 4-hydroxy tamoxifen injections (4-OHT, 75 mg/kg body weight i.p.; Sigma #H6278) to label stress-activated neurons. 4-OHT was prepared the morning of the experiment as follows: 4-OHT was dissolved in 100% ethanol (10% volume of total solution) and added to prepared Captisol solution (360 mg/ml, Research Grade RC-0C7-100), diluted in deionized water (10mg/ml), and homogenized in a water bath with constant sonication (temperature maintained at 42-43 ℃). The LHD1 group received 4-OHT injection 75 minutes after the start of IS1 session, whereas the LHD3 group received 4-OHT injection 75 minutes after the start of ES testing. Both groups were perfused for after 7 days post-LH paradigm (Day 10) to allow time for stable tdTomato expression.

### Timecourse experiment

To further assess the downstream expression pattern of neuronal and extracellular matrix markers and whether perfusion timing and the delay required for tdTomato expression might have affected it, wildtype C57BL/6J mice were perfused at four key timepoints as described in section ii: prior to LH exposure (HC control group), one hour after LH Day 1 training (LH Day 1 group), one hour after LH Day 3 testing (LH Day 3 group) and seven days after LH Day 3 (LH +7 days).

This timeline is illustrated in Figure 4A. The staggered design and the use of wildtype mice allowed us to disentangle the immediate vs. delayed effects of LH stress on PV and PNN expression, independent of TRAP2:Ai14-based genetic labeling.

### Tissue collection and processing

Mice were deeply anesthetized with continuous isoflurane (5%) and transcardial perfusion was performed with 4% PFA solution. Mouse heads were post-fixed for 24 hours at 4°C in a rotary mixer. Sectioning was done using a cryostat (Leica CM3050s) or vibratome (Leica VT1000) at 100 μm thickness. For the mPFC, 3 coronal sections spanning coordinates +1.3 to +1.8 mm relative to bregma were collected. Brain tissue was always preserved using a phosphate-buffered saline (PBS) + 0.1% Azide solution.

A double immunohistochemistry procedure was used to label PV^+^ and PNNs. To label PNNs, we used Wisteria floribunda agglutinin (WFA), a lectin from Japanese wisteria that binds to the CSPGs of PNNs and is a common histochemical PNN marker^42^. Immunohistochemistry incubation was performed at room temperature. First, a blocking solution consisting of 10% Triton X-100, 5% Normal Goat Serum, 2% BSA, and 0.2% Cold Water Fish Serum Gelatin were added to the solution. The next day, a solution consisting of the blocking solution, guineapig anti-PV primary antibody (Synaptic System, cat. #195 308 Guinea pig monoclonal recombinant IgG, 1:2000) and Wisteria Floribunda Lectin Biotinylated (Vector Labs, cat. #B-1355-2, 1:200). Two days later, the brain tissue was washed with 1X PBS five times (10 minutes each). Then, the tissue with incubated with PBS-A with 0.3% Tween-20 with Alexa Fluor 488 AffiniPure Donkey Anti-Guinea Pig IgG (Jackson ImmunoResearch, RRID: AB_2340472, 1:1000) and Alexa Fluor 647-conjugated Streptavidin (Invitrogen, cat. #S21374, 1:3000). Tissues were washed five more times with 1X PBS (10 minutes each) and mounted onto glass slides using ProLong Diamond Antifade Mountant as mounting media.

### Imaging and segmentation analysis

All imaging was performed using a Nikon Eclipse Ti2-E motorized inverted microscope at 10X magnification connected to a Lumencor Aura III Light Engine. Two or three fluorescence channels were captured: PV (emission peak: 488 nm), TRAP+/tdTomato (emission peak: 594 nm), and PNNs (emission peak: 647 nm).

Z-stacks for all three channels were acquired and collapsed using the Enhanced Depth of Focus function in NIS-Elements to generate composite images for quantification of tdTomato, PV, and WFA labeling. TIFF files were generated from raw image data using a custom macro in Fiji/ImageJ to ensure consistent formatting and preprocessing. For downstream analysis, WholeBrain software^43^ implemented in a Virtual Machine Infrastructure (Oracle VM, VirtualBox-7.0.20-163906-Win running Ubuntu OS and RStudio version 1.2 5001), was used to align coronal sections to reference plates from the Allen Mouse Brain Atlas (2011)^44^, enabling unbiased, atlas-guided delineation and quantification of signal intensities from the mPFC subregions. A custom ImageJ macro was used to overlap the extrapolated warped contours of PL and ILA to the correspondent original 16-bit image and the images cropped to reduce the background noise for the following signal segmentation. The same semi-automated Image-J macro was used for preprocessing the images to reduce background noise on each separate channel (background correction via large-sigma Gaussian estimate and iterative Otsu-based thresholding on smoothed background-corrected images), and the generated images were then used for segmentation of TRAP, PV, and PNN channels, that is the detection and quantification of fluorescent puncta, defined as discrete, fluorescently labeled points corresponding to individual neurons or cell body. Specifically, for TRAP and PV signals, a customized python script was used to first denoise & background-correct the images (median filter to remove salt-and-pepper noise), then a duplicate of a heavily blurred version of the image (large-sigma Gaussian) was subtracted to correct for uneven illumination/background across the cropped region of tissue. Then a light Gaussian smoothing, followed by adaptive Otsu thresholding (scaled by a calibrated multiplier) allowed to separate TRAP^+^ or PV^+^ signal from background. Finally, an object filtering based on a calibrated size range (area in µm ²) and pass shape checks (minimum solidity, maximum eccentricity) was used to find soma-shaped objects, excluding noise and non-cell-like blobs. Because WFA signal is characterized by a halo/donut shape signal surrounding the cell bodies, different morphological filters were applied in the python script (fill holes, remove small objects, clear border), and two independent methods were used to avoid false positive (edge glare/gaps). The output for each image consisted in a list of per-object measurements including area, mean intensity (raw and background-corrected), integrated density, eccentricity, centroid x and y. In addition, the area surface of the contour overlapped by the ROIs registration was calculated and later used for density measurements. To evaluate cellular overlap and marker co-expression, the segmented images were binarized and the produced masks overlapped and manually counted. Colocalization was also confirmed in a custom-made python script, by calculating the Euclidean distance (D) between every centroid pair of the two channels of interest, and the puncta pair was marked as colocalized if D was less than a conservative value of 8 µm (based on average PV^+^ diameter and 0.65 µm/pixel resolution). Cell density values were subsequently calculated by normalizing cell counts to the calculated imaged area (mm²) multiplied by 100 µm section thickness to obtain the volume density (mm^3^). Quantitative results represent the summed puncta counts across all layers of the PL and ILA regions divided by the total volume. Colocalization analyses were calculated as the percentage of colocalized cells relative to the total number of cells positive for the reference marker in each region of interest. For the PV and PNN intensity analysis, the mean intensity value in the per-object-measurement list was considered. Because of the inter-animal and region variability, the background corrected value was used.

### Behavioral analysis

Animals’ subjected to LH procedure were classified as being “helpless” (HELP) or “resilient” (RES) based on the most commonly reported indices of helplessness, failures and escape latency, recorded at ES ^41,45,46^. A *k*-means cluster analysis (*k*= 2) was conducted on the database consisting of IS1 and ES group for the TRAPped cohorts, and separately for the LH Day 3 and LH +7 days timecourse cohort ^27,41,45^. Control animals were excluded from the clustering analysis. These behavioral parameters were transformed to *Z* scores, to normalize and prevent differences in the range of the two variables for each cohort that could bias classification in the clustering process. A combined Z-score was calculated as the average sum of the two parameters, with higher values corresponding to more severed helplessness, and used for ANOVA comparison and correlation analysis on groups and phenotypes. Behavioral classification and correlation analysis were conducted in in RStudio (version 2026-05.0+218).

## Statistical Analysis

We used one-way ANOVA analysis to compare phenotypes within each cohort. Repeated mixed effect model or two-way repeated measures ANOVA was conducted to evaluate the effect of two factors and their interaction, such as Timepoint and Phenotype separately for sub-region. To evaluate the effects of sex, stress and timepoint and their interactions, three-way ANOVA tests were conducted separate for each-sub region. Šídák’s or Tukey’s post-hoc tests were performed as recommended to correct for multiple comparisons. Significance was set at α = 0.05. All statistical analyses were conducted in Graphpad Prism 11 or RStudio (version 2026-05.0+218).

## Results

### PL and ILA recruitment during inescapable shock predicts subsequent behavioral outcome

To ask whether prefrontal recruitment during uncontrollable stress relates to later coping behavior in the Learned Helplessness model (LH), we used TRAP2 system to permanently label neurons active during the first inescapable shock session (IS1), before the behavioral phenotype was expressed (Fig. 1A).

In a separate cohort, we also asked whether prefrontal cortex recruitment during the escape session (ES) differed between individuals expressing different coping strategies. Briefly, stressed mice (STR) were subjected to IS during day 1 and day 2 of LH, whereas control mice (CTR) were placed in the shuttle box but no shocks was delivered during the first 2 days. On day 3, all the mice were subjected to a 30-trials escape avoidance test. Individual animals were classified as helpless (HELP) or resilient (RES) based on a 2 clusters k-means cluster analysis computed with escape latency per trial and number of failures (trials in which the animal did not escape from the shock-delivering compartment) (Fig. 1B). The one-way ANOVA conducted to compare the combined behavior z-scores between HELP, RES, and CTR mice in the LH Day 1-cohort, confirmed a main effect of the behavioral group (F_2,14_=26.33, *p<0.0001*), with animals classified as HELP by cluster analysis, having a significantly higher failing rate and average escape latency than both RES and CTR mice (HELPvsCTR: *p<0.0001*; HELPvsRES: *p < 0.00015*) (Fig. 1C). In contrast, RES mice did not differ significantly from CTR mice that never experienced unescapable shocks (*p=0.4622*). Similarly, the LH Day 3-cohort shows a main effect of the behavioral group, with HELP mice having significantly higher combined z-scores than CTR and RES mice, and the last 2 groups being statistically no different (F_2,14_=47.37, *p<0.0001*; post-hoc respectively: HELPvsCTR *p<0.0001*, HELPvsRES *p=0.00000468*, CTRvsRES *p=0.4276*) (Fig. 1C).

Stress exposure alone did not alter level of activation in both PL and ILA subregions, as shown by the two-way ANOVA results for TRAP⁺ count and TRAP^+^ density in either subregion at either timepoint (Fig. 2A; Count: ROIxStressor: F_1,21_=0.2133 *p=0.6490*, ROI: F_1,21_=0.2328 *p=0.6345*, Stressor: F_1,21_=0.4550 *p=0.5073*; Density: ROIxStressor: F_1,21_=0.03675 *p=0.8498*, ROI: F_1,21_=2.131 *p=0.1592*; Stressor, F_1,21_=0.02641 *p=0.8724*). We then investigated whether mice that had shown divergent behavioral coping responses at ES showed any difference in PL and ILA activation levels at IS1, 2 days before expressing the behavioral phenotype, and at ES itself (Fig. 2B). We conducted a between subject mixed-effects analysis to compare TRAP^+^ density among CTR HELP and RES groups in the two independent cohorts IS1 and ES. In PL, we found a main effect of phenotype (F_2,33_=7.332, *p=0.0023*) and timepoint (F_1,33_=23.97, *p<0.0001*), and no significant interaction between the two factors (F_1,33_=2.091, *p=0.1397*) (Fig. 2B). The post-hoc test revealed that RES animals showed higher TRAP⁺ density than HELP animals at IS1 (*p=0.0003*). Importantly, in the independent ES cohort, TRAP⁺ density was lower overall and the phenotype difference was absent. PL activation decreased significantly from IS1 to ES in CTR and RES animals (CTR: *p=0.0345*; RES: *p<0.0001*) (Fig. 2B). In contrast, HELP animals showed relatively low activation at both timepoints, with no significant difference between IS1 and ES. In ILA, only a main effect of the timepoint was found (F_1,33_=12.59, *p=0.0012*), with the post hoc revealing a significant decrease in TRAP-density from IS1 to ES for animals that experienced inescapable shocks (RES: *p=0.0092*; HELP: *p=0.0407*) (Fig. 2B).

**Figure 2:**
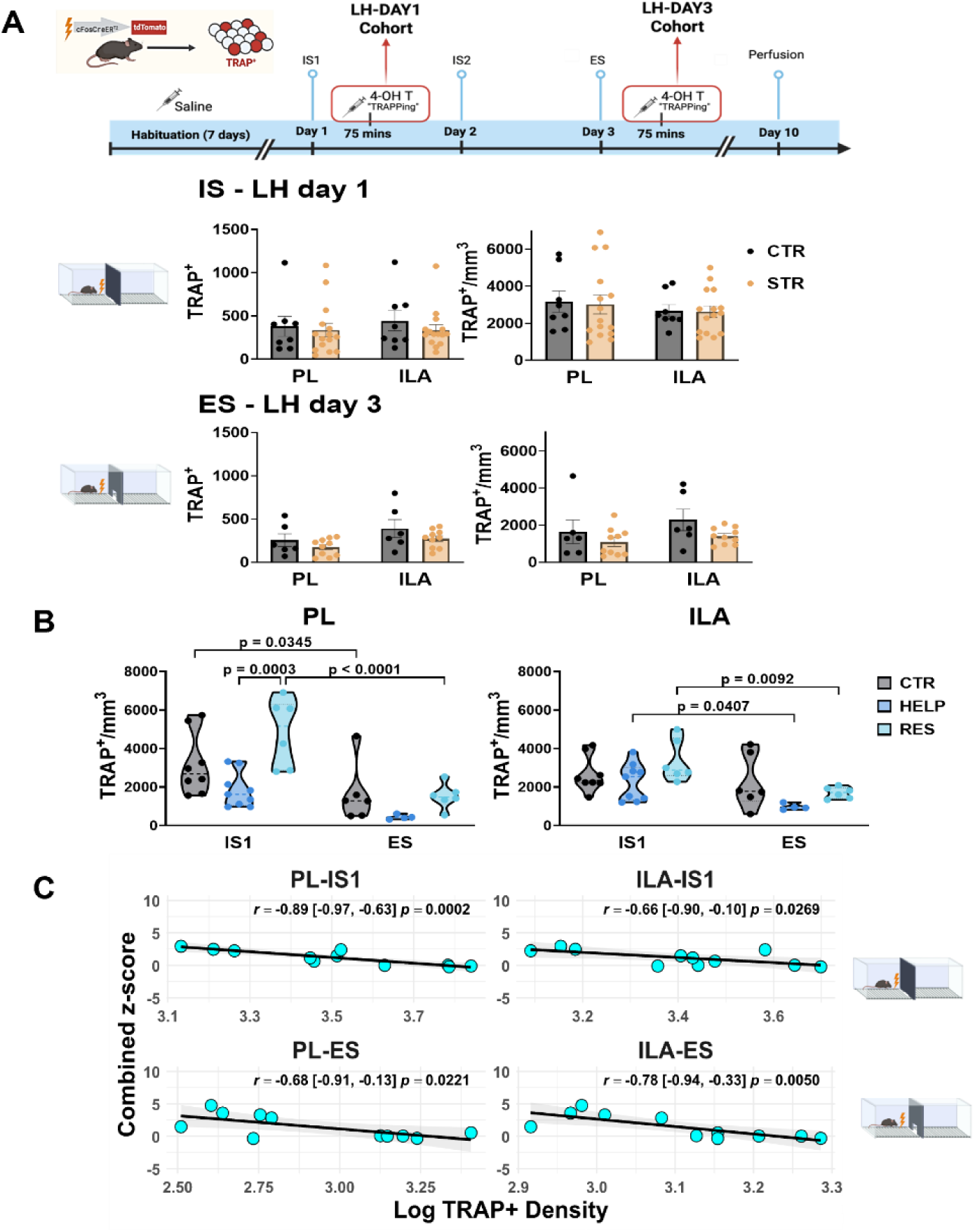
Neuronal activation of mPFC at day 1 and day 3 of LH. (A) Experimental design for IS1-LH-Day 1 cohort) and ES-LH-Day 3 cohort. Mice received 4-OHT 75 minutes after IS1 or ES and were perfused on day 10. TRAP+ cell counts and densities were quantified in the prelimbic (PL) and infralimbic (ILA) subregions. Control (CTR) and shocked (STR) animals were analyzed. Bars represent the mean ± SEM, and points represent individual animals. The effects of stress exposure and region were evaluated using two-way ANOVA with stress and ROI as factors. (B) TRAP^+^ cell density in PL and ILA per behavioral group and subregion. Shocked animals were classified as helpless (HELP) or resilient (RES) based on their escape behavior. IS1 and ES were assessed in independent cohorts; therefore, comparisons between sessions represent between-animal rather than within-animal differences. Violin plots show the distribution of individual-animal values, with horizontal lines indicating the median. Data were analyzed separately for PL and ILA using mixed-effects models with behavioral phenotype and labeling session as factors, followed by post hoc comparisons adjusted for multiple testing. **(C)** Correlation between log-transformed TRAP* density and the combined behavioral z-score in shocked animals. Higher behavioral z-scores indicate longer escape latencies and more escape failures. Points represent individual animals, solid lines show linear regression fits, and shaded regions show 95% confidence intervals. Correlations were evaluated using two-sided Pearson tests.

Because IS1 and ES TRAP-labelling was performed in separate cohorts, a within-animal trajectory was not possible and comparisons were interpreted as two independent photographs of neuronal activation at the two LH timepoints investigated. The results indicated that the increased level of activation via TRAP proxy reflects elevated PL recruitment in resilient animals in presence of stress, and a deficit in helpless animals to recruit the subregion. In addition, the CTR group had highly PL activation at first exposure to the shuttle box at IS1, independently from the shocks, not significantly different than RES mice in escapable conditions. Because the TRAP2 system leads to the Cre-dependent expression of tdTomato from an activity-dependent promoter, the amount of labelled neurons reflects a robust activity beyond the FOS expression. Therefore, the overall reduction in TRAP density from LH Day 1 to LH Day 3, in both PL and ILA subregions, could reflect the change in duration and amount of shocks between the two sessions for the stressed groups.

LH behavior is classically based on the binary classification of phenotypes in resilient and helpless groups. Still, the behavioral analysis revealed a continuous distribution of the number of failures to escape and average latencies to escape (Fig. 1B). In order to investigate whether PFC activation correlates to coping responses independently of the phenotype classification, we calculated a combined behavioral z-score of failures and latency parameters, with higher z-score corresponding to reduced resilience/increase helplessness. Consistent with the group-level analysis, we found that TRAP⁺ density during IS1 was inversely related to the combined behavioral z-score in shocked animals, strongly for PL than ILA (Pearson r_PL_ = −0.89, 95% CI [-0.97,-0.63], *p=0.0002*; r_ILA_ = −0.66, 95% CI [-0.90,-0.10], *p=0.0269*) (Fig. 2C). The relationship was preserved in the independent ES cohort, although the strength of the correlation was higher for ILA than PL (r_PL_= −0.68, 95% CI [-0.91,-0.13], *p=0.0221*; r_ILA_= −0.78, 95% CI [-0.94,-0.33], *p=0.0050*). Thus, prefrontal recruitment measured during the first uncontrollable stress session, before behavioral divergence, scales continuously with subsequent escape performance.

### PV expression tracks behavioral outcome rather than stress exposure

To investigate how the observed changes in neuronal activation were associated with LH behavior in the two mPFC subregions, both cohorts of mice were perfused one week after ES session to permit the full expression of the tdTomato labelling, and the brains processed for immunostaining procedures (Fig. 3A). Because PV⁺ play a pivotal role in regulating local circuit activity within the mPFC, we first questioned whether LH was affecting PV expression in PL and ILA. The expression of PV protein and its gene, *Pvalb*, is tightly coupled to the level of neuronal electrical and synaptic activity in the brain ^46^. High network activity or sensory enrichment increases PV expression, whereas sensory deprivation or reduced firing decreases PV levels ^47,48^. Thus, we quantified PV⁺ interneurons in PL and ILA one week after LH (Fig. 3B). We first examined the effect of stress on the PV⁺ interneurons count normalized by volumes (PV^+^/mm^3^) in the two subregions of PFC (Fig. 3B). The two-way ANOVA revealed a main effect of ROI (F₁, ₂₅ = 5.297, p = 0.030), indicating that PV⁺ interneuron density differed overall between PL and ILA. However, there was no significant main effect of stress and no significant stress × ROI interaction, indicating that PV⁺ density did not differ between CTR and STR animals in either subregion (Fig. 3B). When shocked animals were stratified by phenotype observed at ES (Fig. 3C), a main effect of both phenotype and ROI was found (Phenotype: F_1,24_=5.908 *p=0.023*, ROI: F_2,24_=5.658 *p=0.010*), but no interaction between the two variables. The post hoc revealed RES (p = 0.0037) (Fig. 3B, right). A similar but nonsignificant pattern was observed in ILA. RES did not differ from CTR in either subregion. These findings suggest that PV⁺ density is associated with behavioral outcomes rather than stress exposure alone.

**Figure 3:**
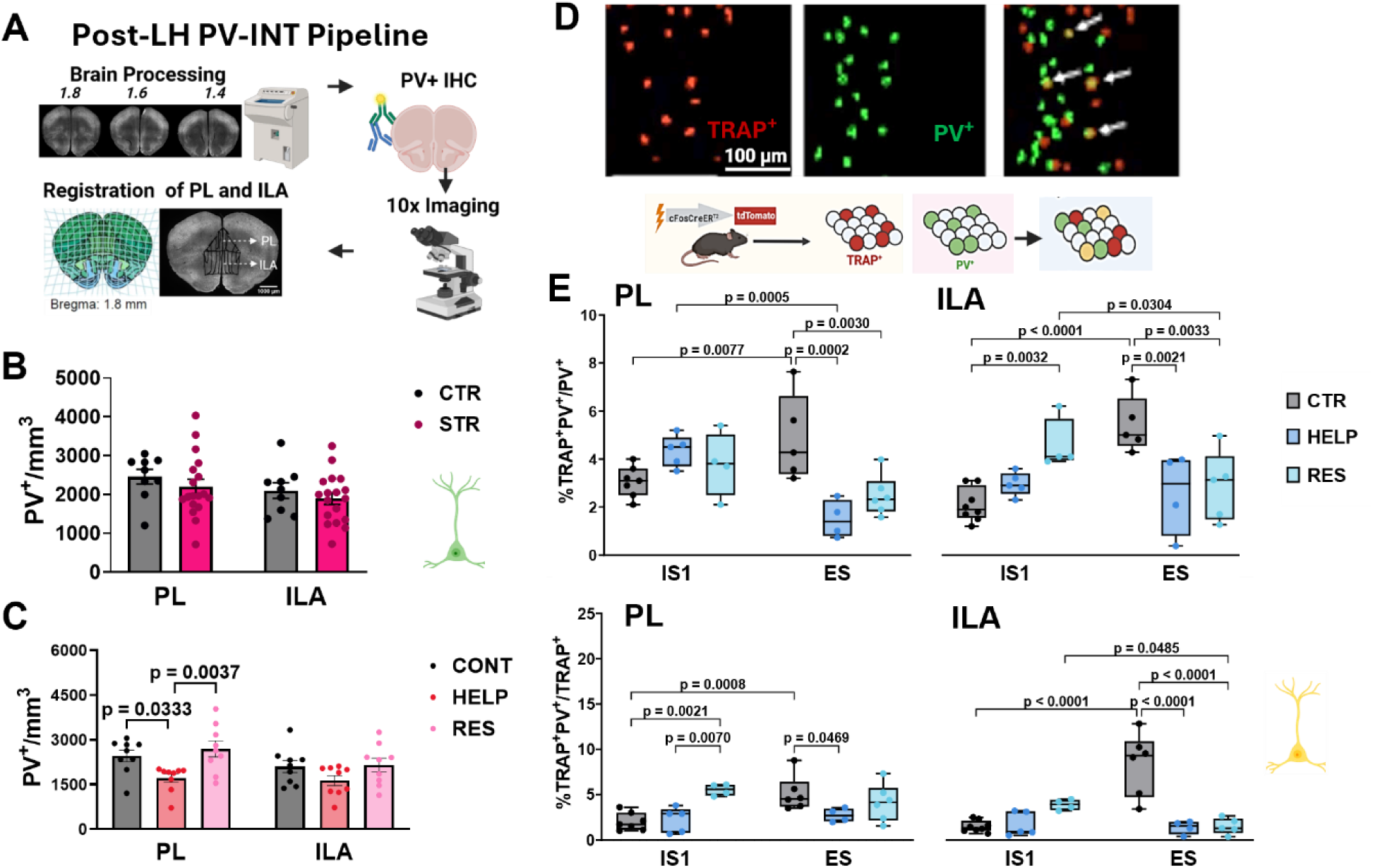
PV expression following LH paradigm. **(A)** Tissue-processing workflow used to quantify PV^+^ interneurons in the PL and ILA subregions. Coronal sections spanning approximately 1.4-1.8 mm anterior to bregma were immunostained for PV and imaged at 10* magnification. **(B)** PV^+^ interneuron density in CRTL and STR animals seven days after the escape test, with shocked animals pooled across behavioral phenotypes. Bars represent the mean ± SEM, and points represent individual animals. Data were analyzed using two-way ANOVA with stress exposure and ROI as factors. Sample sizes: CTR, n = 9; STR, n = 18. **(C)** PV^+^ interneuron density after shocked animals were stratified as HELP or resilient RES. Bars represent the mean ± SEM, and points represent individual animals. Data were analyzed using two-way ANOVA with behavioral phenotype and ROI as factors, followed by post hoc comparisons adjusted for multiple testing. Sample sizes: CTR, n = 9; HELP, n = 7; RES, n = 11. **(D)** Representative images of TRAP^+^ neurons, PV^+^ interneurons, and TRAP-PV colocalization. Arrows indicate TRAP^+^PV^+^ cells. Scale bar, 100 pm. **(E)** Recruitment of PV^+^ interneurons during IS1 and ES in independent cohorts. Upper plots show the percentage of PV^+^ interneurons that were TRAP^+^ (TRAP^+^PV^+^/PV^+^), whereas lower plots show the percentage of TRAP^+^ neurons that were PV^+^ (TRAP^+^PV^+^/TRAP^+^). Boxes indicate the median and interquartile range, whiskers indicate the minimum and maximum values, and points represent individual animals. Data were analyzed using between-subject two-way ANOVA with behavioral phenotype and labeling session as factors, followed by post hoc comparisons adjusted for multiple testing. Exact p values are shown. Sample sizes: IS1 cohort: CTR, n = 6; HELP, n = 5; RES, n = 4; ES cohort: CTR, n = 5; HELP, n = 5; RES, n = 5.

**Figure 4:**
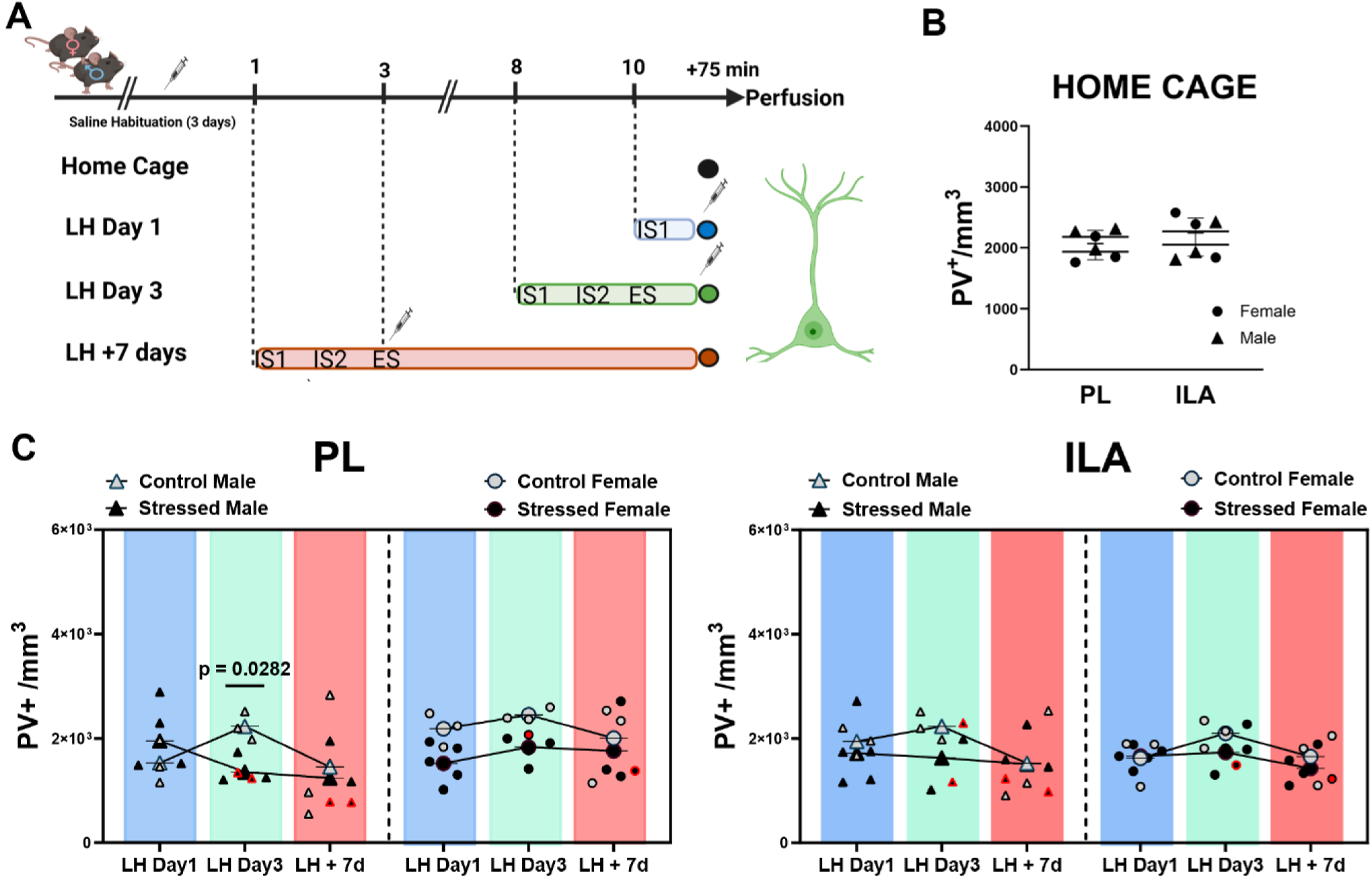
PV expression during LH timecourse. **(A)** Staggered endpoint experimental design. Separate cohorts were perfused 75 minutes after the first inescapable-shock session (LH Day 1), 75 minutes after the day 3 escape test (LH Day 3), or seven days after the escape test (LH +7 days). The LH +7 days endpoint matched the time point examined in the preceding experiments. An unmanipulated home-cage group provided a reference baseline. **(B)** Baseline PV^+^ interneuron density in PL and ILA of home-cage mice. Circles denote females and triangles denote males. Horizontal lines and error bars represent the mean ± SEM. Each point represents one animal. Sample size: n = 7 mice (3 females and 4 males). **(C)** PV^+^ interneuron density in PL and ILA according to endpoint, stress exposure, and sex. Open symbols denote control animals, filled symbols denote shocked animals, circles denote females, and triangles denote males. Large symbols and connecting lines represent group means; error bars show the SEM. Red-outlined symbols identify shocked animals classified as “helpless” and are shown for descriptive purposes only because the LH Day 3 and LH +7 days cohorts contained only two helpless mice per endpoint. Data were analyzed separately for PL and ILA using three-way ANOVA with sex, stress exposure, and endpoint as factors, followed by post hoc comparisons adjusted for multiple testing.

### PV recruitment is outcome-related during inescapable shock and exposure-related during escape testing

We further ask whether PV interneurons were themselves recruited during LH, at the two timepoints investigated. Therefore, we quantified TRAP–PV co-localization in the two cohorts IS1 and ES (Fig. 3D). In both subregions, the between-subject two-way ANOVA revealed a significant interaction between Phenotype and Timepoint (PL: F_2,26_= 6.690, *p=0.0045*; ILA: F_2,26_= 23.39, *p<0.0001*), and a main effect of the Phenotype (PL: F_2,26_= 5.868, *p=0.0079*; ILA: F_2,26_= 11.52, *p=0.0003*) (Fig. 3E top). The post-hoc analysis adjusted for multiple comparisons indicated that the proportion of TRAP⁺ neurons expressing PV was higher in the RES mice than in CTR and HELP mice at IS1, significant only in PL (RES vs CTR *p=0.0021*; RES vs HELP *p=0.0070*). In the ES cohort, CTR animals showed a marked increase in the PV⁺ fraction of the activated ensemble in ILA relative to both shocked groups (CTR vs HELP *p<0.0001*; CTR vs RES *p<0.0001*), and a slight increase in PL relative to HELP (*p=0.0469*), while HELP and RES were indistinguishable in both subregions. Notably, RES at IS1 and CTR at ES exhibited similar proportions of PV⁺ neurons within the activated prefrontal ensemble; in both cases, the animals were experiencing foot shock for the first time.

Because the previous measurements of PV^+^ interneurons recruitment were dependent on the total TRAP⁺ density being the denominator, which itself differed between cohorts (Fig. 2B), we also calculated a reciprocal measure: the percentage of PV^+^ interneurons that were TRAP⁺ in each prefrontal subregion at both of timepoints (Fig. 3E bottom). The two-way ANOVA showed again a significant interaction between Phenotype and Timepoint in both subregions (PL: F_2,26_= 12.95, *p=0.0001*; ILA: F_2,26_= 15.04, *p<0.0001*). At IS1, the fraction of activated PV^+^ among the total number of PV^+^ were similar between groups in PL, whereas it was higher for RES compared to CTR in ILA (*p=0.0032*). AT ES, CTR showed a higher fraction of activated PV^+^ than the other two groups in both subregions (PL: CTR vs HELP *p=0.0002*, CTR vs RES *p=0.003*; ILA: CTR vs HELP *p=0.0021*, CTR vs RES *p=0.0033*). Across cohorts, the proportion of PV+ interneurons recruited was generally lower at ES than at IS1. This difference was significant for CTR and HELP in PL (p=0.0077 and p=0.0005 respectively), and for CTR and RES in ILA (p<0.0001 and p=0.00304 respectively). Once again, normalizing the number of PV⁺ recruited to the more stable total PV⁺ population (**Sup. Fig. X),** shows the same stress exposure driven at ES in both prefrontal subregions, suggesting that the result was unlikely to be driven by denominator variation alone. At IS1, the results were again consistent across the two measurements in ILA where RES mice had a modestly higher proportion of activated PV⁺ interneurons than CTR mice. A similar trend was observed in PL only when recruited PV⁺ interneurons were normalized to the total number of activated neurons.

Together these data indicate that PV recruitment during uncontrollable stress differs by subsequent behavioral outcome, whereas during the escape test it differs by prior stress exposure irrespective of outcome — the same temporal signature seen for overall prefrontal recruitment.

### A stress effect on PV expression emerges at the escape test and is no longer detectable one week later

The PV difference observed between behavioral phenotypes on day 10 (Fig. 3C) together with the dynamical recruitment of PV interneurons across LH sessions (Fig. 3E), raised the question of when PV protein expression first changes relative to stress exposure. To address this, we used a staggered design in which separate cohorts underwent the LH paradigm to different endpoints: 15 min after the first inescapable shock session (LH Day 1), 15 min after the day 3 escape test (LH Day 3), or seven days after the escape test (LH +7 days). The LH +7 days endpoint matched the time point examined in the preceding experiments. All animals were perfused on the same day and processed in a single immunohistochemical batch, with slides from different groups interleaved during processing (Fig. 4A). This design matches tissue age, staining batch, and imaging conditions across timepoints. An unmanipulated home-cage group provided a reference baseline (Fig. 4B). Control animals in each timepoint cohort received equivalent chamber exposure, handling stress mice, with the exception of shock-free exposure at IS1 and IS2. We first conducted a three-way ANOVA to see the effect of sex, stress, and timepoint separately for the two subregions. In PL, we found a significant interaction between sex and stress on PV⁺ density (PL: F_1,18_= 10.51, *p=0.0045*) (Fig. 4C left). The multiple comparative analysis showed no differences between shocked and control animals after the first inescapable shock session or seven days after LH completion in either subregion.

In contrast, at LH Day 3 when the escape test took place, PV⁺ density was lower in shocked animals than in controls, with the difference reaching significance in males (*p=0.0282*). No significant main effects or interactions were detected in ILA, although the pattern was similar to that observed in PL, with control animals having a higher mean PV⁺ density than shocked animals shortly after ES (Fig. 4C left). Because each timepoint was sampled in a separate cohort, these comparisons are between-animal and are interpreted as independent snapshots rather than a within-animal trajectory. In summary, after IS1 and seven days after the escape test, no group difference was detectable in either subregion. The absence of a shocked-versus-control difference at LH +7 days was consistent with the result obtained at the equivalent endpoint in the independent main cohorts (Fig. 3B, left). Together, these findings suggest that the PV⁺ density reduction in stressed animals observed at LH Day 3 was transient and was no longer detectable seven days after the escape test, indicating resolution of an earlier effect rather than absence of any stress effect on PV expression. In addition, the temporal pattern of PV expression therefore did not differ between sexes, though the effect of the stressor was more marked in males.

Animals in the LH Day 3 and LH +7 days cohorts completed the full paradigm and could be classified as helpless or resilient. These cohorts were not powered for phenotype-level comparison (n=2 helpless per timepoint) and no statistical tests were performed on the behavioral outcomes. Therefore; helpless animals were indicated in Fig. 4C for descriptive purposes only (red border points). Across sexes and subregions, helpless animals were occupying the lowest values of the shocked group seven days later while resilient animals spanned the control range. This descriptive pattern is consistent with the phenotype difference established on Day 10 (LH +7 days) in an independently powered cohort (Fig. 3C). The trend is already visible at LH Day 3 and suggests that the reduction in PV density at the escape test is shared across shocked animals, whereas its persistence is restricted to those expressing helpless behavior.

### PV, not PNN, tracks stress exposure, with the two markers becoming coupled selectively in ILA

Mature perineuronal nets (PNNs) increase PV^+^ interneuron–mediated inhibition by inhibitory synapses originating from PV^+^ interneurons and stabilizing excitatory synapses onto PV^+^ interneurons. Having shown that PV density changes progressively with stress exposure, decreasing at the escape test and staying low in HELP animals one week after LH, we next asked whether PNN expression visualized by Wisteria floribunda agglutinin (WFA) follows a comparable trajectory in the same timecourse cohorts (Fig. 5A-B, left panels). In contrast to PV, PNN intensity showed no significant stress effect across the timecourse in either subregion, indicating that PNN intensity alone is not a robust marker of stress exposure over this time windows. PV intensity, by contrast, showed significant stress effects in both subregions (Fig. 5A-B, right panels) (PL: Stress F_2,31_= 4.983, *p=0.013*; ILA: Stress F_2,31_= 4.306, *p=0.022*), and a marked significant timepoint effect in ILA (Timepoint F_1,31_=8.083, *p =0.008*). The post hoc revealed a significantly higher PV intensity in ILA at LH Day 3 (*p=0.014*). Those results are consistent with a progressive structural and functional change in the PV interneuron population in PL and ILA subregions following the inescapable shocks experienced during the LH paradigm.

**Figure 5:**
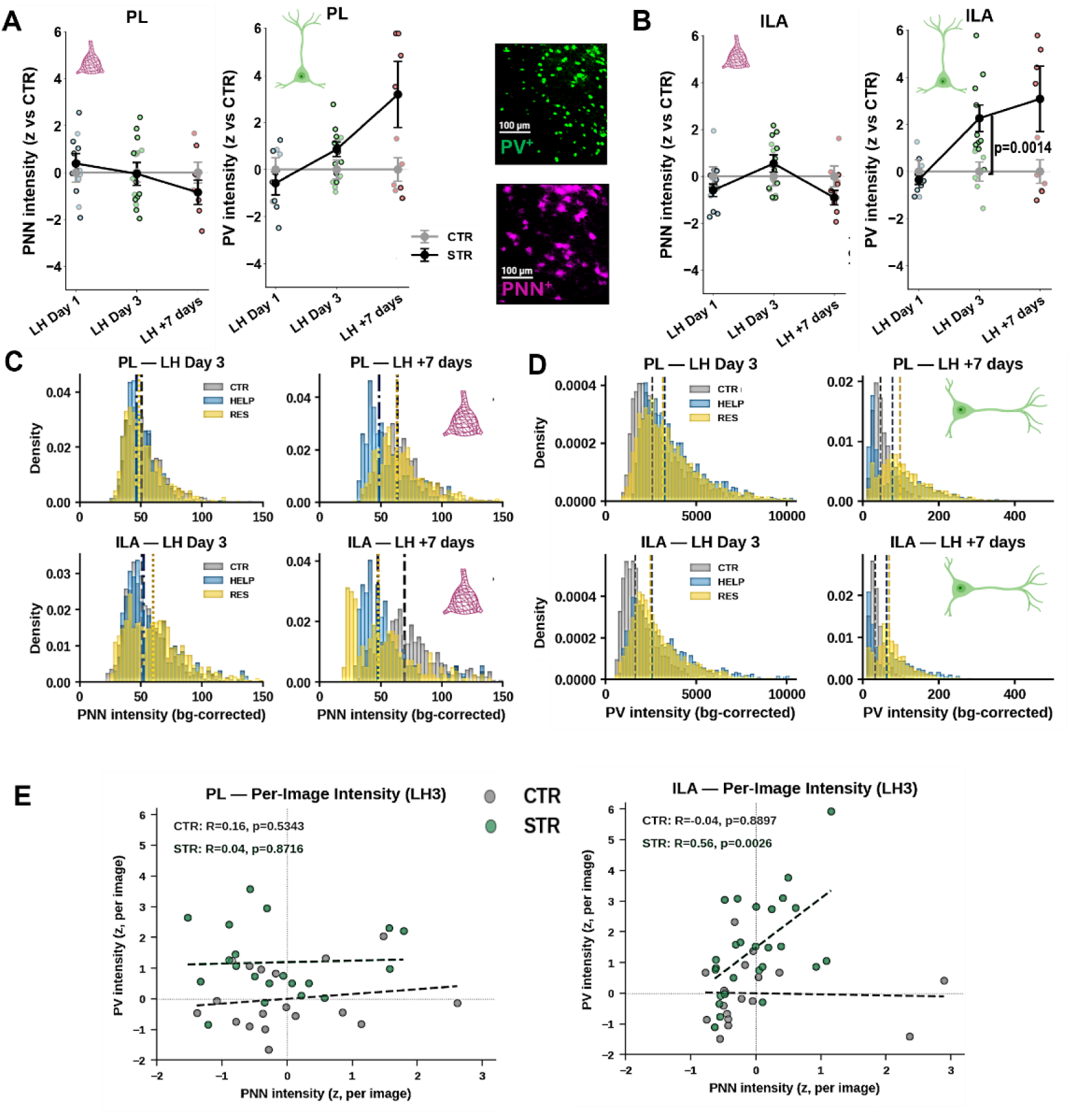
PV intensity, PNN intensity, and their relationship across LH timecourse. Standardized perineuronal-net (PNN) intensity, measured using WFA labeling, and PV fluorescence intensity in PL **(A)** and ILA(B) at LH Day 1, LH Day 3, and seven days after the escape test. Values are expressed as z-scores relative to the corresponding control group. Single points represent individual animals, and large symbols and error bars represent the mean ± SEM. Representative PV and PNN fluorescence images are shown in A on the right. Data were analyzed separately by region using two-way ANOVA with stress exposure and timepoint as factors, followed by post hoc comparisons adjusted for multiple comparisons. **(C-D)** Distributions of background-corrected PNN intensity **(C)** and PV intensity **(D)** in CTR, HELP, and RES groups at LH Day 3 and LH +7 days, shown separately for PL and ILA. Dashed vertical lines indicate the group medians. Histograms represent measurements from individual PNN-associated PV^+^ cells pooled across animals. (E) Correlation between PNN and PV intensity at LH Day 3 in PL and ILA. Points represent individual images from CTR or STR animals, and dashed lines show group-specific linear fits. Correlations were evaluated using two-sided Pearson tests.

When we examined the full intensity distributions per counted object across the behavioral phenotypes (HELP VS RES) for LH Day 3 and LH +7 days groups (Fig. 5C-D), PNN intensity distributions remain largely overlapping across groups at each timepoint (Fig. 5C), while PV intensity distributions show a rightward shift with a stress effect that becomes more pronounced at later timepoints and separates HELP from RES animals by LH +7 days (Fig. 5D), reinforcing that the stress signature in this dataset is driven more clearly by PV expression than by PNN intensity itself.

Given that PNN intensity alone did not track stress exposure, we asked whether PNN and PV levels were instead coupled to one another within stressed animals (Fig. 5E). At LH Day 3, PNN and PV intensity in PL were uncorrelated regardless of inescapable stress exposure, indicating no PNN-PV relationship in this subregion at the time window investigated (Fig. 5E top). In ILA, however, PNN and PV intensity were significantly correlated in STR animals (R=0.56, p =*0.0026*) but not in CTR (R =-0.04, p =0.8897) (Fig. 5E bottom). This dissociation suggests that stress exposure induces a PNN-PV coupling that is specific to ILA and specific to the stressed state, rather than reflecting a constitutive relationship between the two markers present in unstressed conditions.

## Discussion

The present findings indicate that vulnerability to helplessness is reflected in prefrontal recruitment before being behaviorally expressed. We found that the density of TRAP⁺-activated neurons in prelimbic cortex (PL) during the first inescapable shock session was inversely related to the combined behavioral z-score measured at the escape test two days later, and resilient animals showed higher PL recruitment than helpless animals at this early timepoint. Prefrontal engagement therefore precedes, rather than follows, the coping strategy the animal will express, consistent with the view that variability in corticolimbic circuits predicts divergent responses to stress rather than resulting from them ^17,49^. Notably, control animals encountering the shuttle box for the first time, in the absence of any shock, activated PL to a level comparable to that of resilient animals exposed to inescapable shocks, whereas helpless animals failed to reach this level in either inescapable or escapable conditions. PL activation in this paradigm may thus reflect a general signal of coping readiness rather than a response specific to shock delivery, with helplessness corresponding to a failure to engage it. This interpretation is consistent with prior evidence that PL is preferentially activated by behavioral control over a stressor rather than by the stressor itself, that such control-related engagement can be elicited by a novel controllable context, and that prior experience of control can protect against the effects of subsequent uncontrollable stress ^50–52^.

In ILA, by contrast, activation declined uniformly from IS1 to ES in shocked animals, without distinguishing coping phenotype. This pattern is consistent with habituation to repeated homotypic stress, whereby stress-induced activation of limbic and prefrontal regions decreases upon re-exposure to the same stressor irrespective of coping strategy^53,54^. Because ventral mPFC activity has been implicated as necessary for the development of such habituation rather than as a passive consequence of it ^53^, ILA engagement during this window may reflect cumulative stress exposure, and potentially contribute to habituation, rather than the individual differences in coping that characterize PL. In a recent paper, Starsved and colleagues reported the opposite regional dissociation when comparing acute and chronic restraint stress, in which prelimbic cortex showed partial habituation following repeated stress, whereas infralimbic and orbitofrontal responses were unaffected by chronicity, with infralimbic habituation instead emerging only in females in low-estradiol phases ^55^. This discrepancy may reflect differences between a two-session inescapable-shock protocol and multi-week restraint regimens, or between foot shock and restraint as stressors, and warrants further investigation.

A comparable temporal pattern was observed at the level of PV⁺ interneuron recruitment during LH in the two prefrontal regions. During the first inescapable shock, resilient animals recruited a larger PV⁺ interneuron fraction of the activated PL and ILA ensembles than helpless or control animals, indicating that the phenotype-predictive signal reflects preferential engagement of inhibitory neurons rather than a general increase in activation. During the escape test, the pattern was instead associated with prior shock exposure. control animals, which experienced the stressor for the first time in an escapable condition, recruited the largest PV⁺ fraction in both subregions, whereas helpless and resilient animals, both unescapable shock-experienced by this point, did not differ from one another. PV⁺ interneuron recruitment

Importantly, we found that one week after LH, PV⁺ density did not differ between shocked and control animals when behavioral phenotypes were pooled. However, phenotype - stratified analysis showed lower PV⁺ density in the PL of helpless mice than in control and resilient mice, with a similar nonsignificant pattern in ILA. We reasoned that a structural difference in terms of numbers of this subpopulation of interneurons between the groups was unlikely to be the cause, but rather reflecting a functional dynamic reorganization of the activity-dependent parvalbumin protein expression. Indeed, PV⁺ interneurons play a key role in activity-dependent neuronal plasticity, a fundamental mechanism through which the nervous system adapts to sensory experience. Parvalbumin itself is a calcium-binding protein that acts as a slow intracellular calcium buffer to regulate short-term synaptic plasticity in these fast-spiking cells. Our timecourse staggered design experiment aimed to track the temporal pattern of the changes observed one week after LH. We found that, after the first inescapable shock, PV⁺ density was reduced in shocked animals at the escape test and no longer differed between groups seven days later when animals were pooled across phenotypes. When this later timepoint was stratified by phenotype, helpless animals occupied the lowest values of the shocked distribution, a pattern already apparent at Day 3. Together, these results suggest that inescapable stress produces a transient, exposure-wide suppression of PV⁺ expression around the time of the escape test, from which most animals recover over the following week; the failure to recover, rather than the initial reduction, distinguishes the helpless phenotype. Finally, the sex-by-stress interaction on PL PV⁺ density, together with the significant shocked-versus-control difference observed in males, suggests that the magnitude of this effect may be sex-dependent, although the small size of the staggered cohorts is not conclusive.

In contrast to PV⁺ cell density, PV⁺ intensity, reflecting protein expression per cell, showed a distinct pattern. Pooled across the staggered cohorts, PV⁺ intensity increased with stress in both PL and ILA, most markedly in ILA at Day 3. When animals were stratified by phenotype at Day 3 and LH +7 days, this stress-related increase resolved into a phenotype effect, with helpless animals occupying the right-shifted, higher-intensity portion of the distribution relative to resilient animals and the separation increasing between the two timepoints. These findings are not inconsistent with the density results; rather, the two measures may reflect complementary aspects of the same reorganization, whereby fewer PV⁺ cells are detectable in the shocked population overall, while the remaining cells, particularly in animals that become helpless, upregulate PV expression. This pattern accords with an over-inhibition account of stress-induced prefrontal dysfunction, in which chronic stress increases both the activity and the expression of prefrontal PV⁺ interneurons, producing a more strongly inhibited network and associated anxiety-and depression-like behavior ^56^. Chemogenetic inhibition of infralimbic PV⁺ interneuron activity attenuates the behavioral and physiological consequences of chronic stress ^35^, raising the possibility that the elevated PV⁺ intensity observed in helpless animals by LH +7 days contributes to maintaining the phenotype rather than serving merely as a correlate of it, a possibility that could be tested directly using a comparable approach in this paradigm.

Activity-dependent plasticity in PV⁺ cells is gated by remodeling the extracellular perineuronal nets. Therefore, we hypothesized that changes in PNN expression might precede or follow PV⁺ interneuron dynamics. In contrast, we found that PNN did not parallel the changes observed for PV in the same timecourse experiment. WFA intensity did not differ significantly with stress exposure in either subregion across the time course and remained largely overlapping across phenotypes even at LH +7 days, when PV distributions most clearly separated helpless from resilient animals. Nonetheless, PNNs were not without a stress-related signature: PNN and PV intensity were correlated specifically in ILA of stressed animals, with no such relationship in unstressed controls or in PL under either condition. Because this coupling was present only in the stressed state and only in ILA, it more likely reflects a stress-induced reorganization of the local relationship between the two markers than a constitutive anatomical association between PV⁺ cells and their surrounding nets.

Across all measures, PL differences were most prominent early, during the uncontrollable stressor, whereas the ILA effects, including the elevated PV⁺ recruitment in controls at the escape test and the stress-selective PNN-PV coupling, emerged later, once escape was possible. This temporal dissociation is consistent with proposed roles for PL in the appraisal of behavioral control and for ILA in the expression and stabilization of an adaptive response once a coping strategy is available ^49,51,56^. Importantly, Wallsten et al. recently found that escapable stress selectively increased recruitment of PV/PNN-associated neurons in the rat IL, whereas both escapable and inescapable stress increased c-Fos intensity in PV/PNN cells across PL and IL^58^. Our findings complement this work by comparing PV-interneuron recruitment across distinct stages of learned helplessness and behavioral phenotypes, and by identifying a stress-associated relationship between PV and PNN intensity in ILA despite the absence of a uniform change in PNN intensity.

The different temporal patterns of PV and PNN measures further refine this distinction. PV⁺ interneuron recruitment and expression changed across the transition from uncontrollable stress to escape testing, whereas PNN intensity remained comparatively stable. PNN remodeling therefore does not appear to be required for the early PV-related differences associated with subsequent coping behavior, at least as measured by WFA intensity at the endpoints examined here. Instead, the stress-specific association between PNN and PV intensity in ILA suggests that changes in their local relationship may emerge later, after repeated stress exposure. These findings identify PV⁺ interneurons as a more sensitive marker of the transition between vulnerability and behavioral expression, while suggesting that PNN involvement may occur through altered coupling with PV rather than through a uniform change in PNN intensity.

Several features of the design constrain interpretation. IS1 and ES recruitment were assessed in independent cohorts, so comparisons across timepoints represent between - subject snapshots rather than within-animal trajectories. The staggered LH Day 3 and LH +7 days cohorts were not powered for phenotype-level comparisons (n = 2 helpless animals per timepoint), and the observation that helpless animals occupied the lowest values of the shocked distribution is therefore descriptive rather than confirmatory. The PNN-PV correlation in ILA, although robust, cannot establish the direction of the relationship or whether both measures reflect a common upstream signal.

Taken together, these findings indicate that vulnerability to helplessness begins to emerge in the prefrontal cortex before the animal fails to escape. PV⁺ interneuron dynamics, more than PNN structure, tracked the transition from early stress processing to later expressed coping behavior. The stress-dependent PNN-PV coupling observed in ILA identifies a distinct, later stage of circuit reorganization and suggests that the molecular and cellular processes associated with initial vulnerability may differ from those accompanying the subsequent expression or maintenance of helpless behavior. Distinguishing these stages may help identify neural mechanisms that promote adaptation to adversity.

## Author Contribution

L.A. conducted experiments, analyzed data, contributed to the study design, and wrote the manuscript. S.W. and M.A. performed experiments. A.M. analyzed data, designed and supervised the study, and wrote the manuscript. D.D. contribute to the design of the study.

## Acknowledgement

We thank Dr. Benedetta Di Cesare for assistance with the immunohistochemistry protocol development and Dr. Giulia Zanni for valuable discussion. Panel A in Figures 1-4 were created with BioRender.com.

## Supplementary Figures

**Figure S1:**
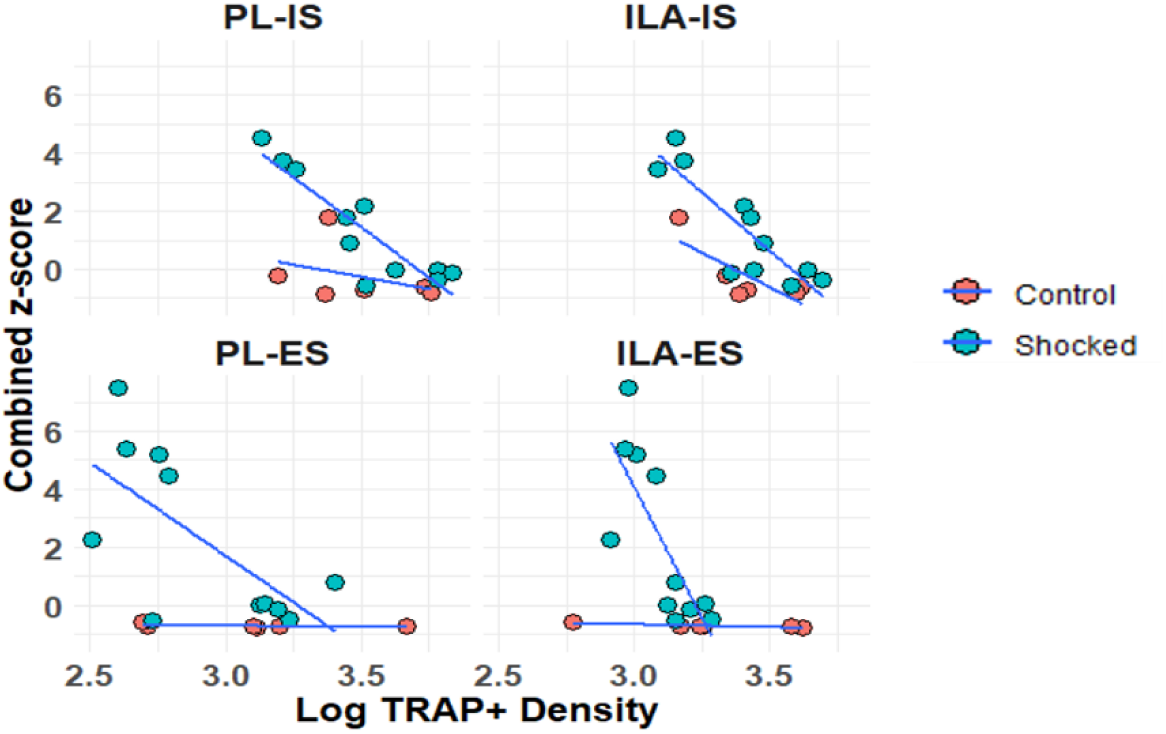
Behavioral correlates of TRAP activation in mPFC. Correlation between log-transformed TRAP^+^ density and the combined behavioral z-score in control (pink) and shocked (turquoise) animals. Higher behavioral z-scores indicate longer escape latencies and more escape failures. Points represent individual animals; solid lines show linear regression fits.

**Figure S2:**
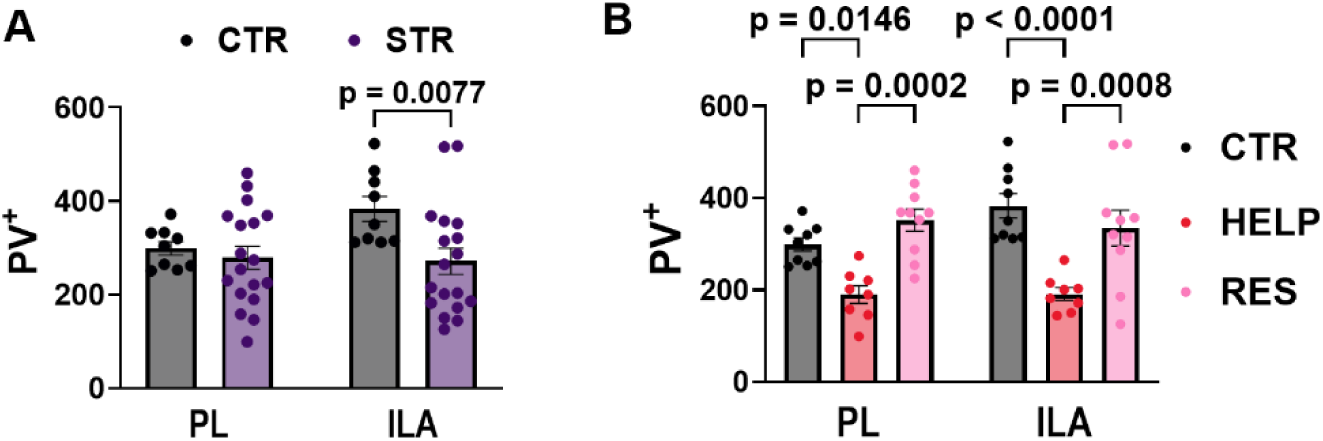
PV interneuron counts following LH. **(A)** PV^+^ interneuron count in CRTL and STR animals seven days after the escape test, with shocked animals pooled across behavioral phenotypes. **(B)** Same of A but STR mice were separated in HELP and RES animals. Bars represent the mean ± SEM, and points represent individual animals. Data were analyzed using two-way ANOVA with stress exposure and ROI as factors.

**Figure S3:**
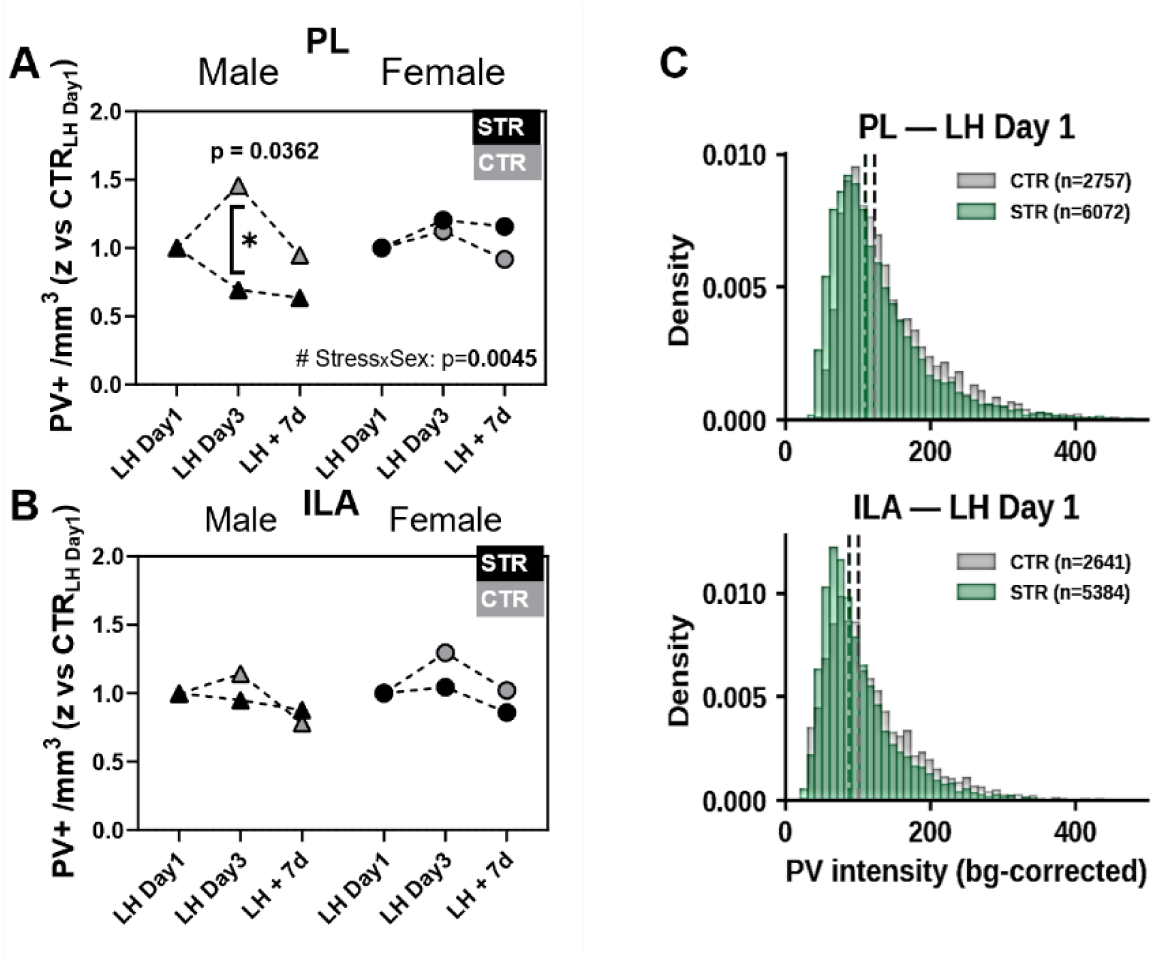
PV+ density across cohort across LH timecourse. Standardized PV+ density in PL **(A)** and ILA **(B)** at LH Day 1, LH Day 3, and seven days after the escape test. Values are expressed as z-scores relative to the corresponding control group. Large symbols and error bars represent the mean ± SEM, and the p-value reported corresponds to post-hoc following three-way ANOVA. **(C)** Distributions of background-corrected PV intensity in CTR, STR groups at LH Day 1, shown separately for PL and ILA. Phenotypes classification not present because endpoint preceding ES testing. Dashed vertical lines indicate the group medians. Histograms represent measurements from individual PV^+^ cells pooled across animals.

**Figure S4:**
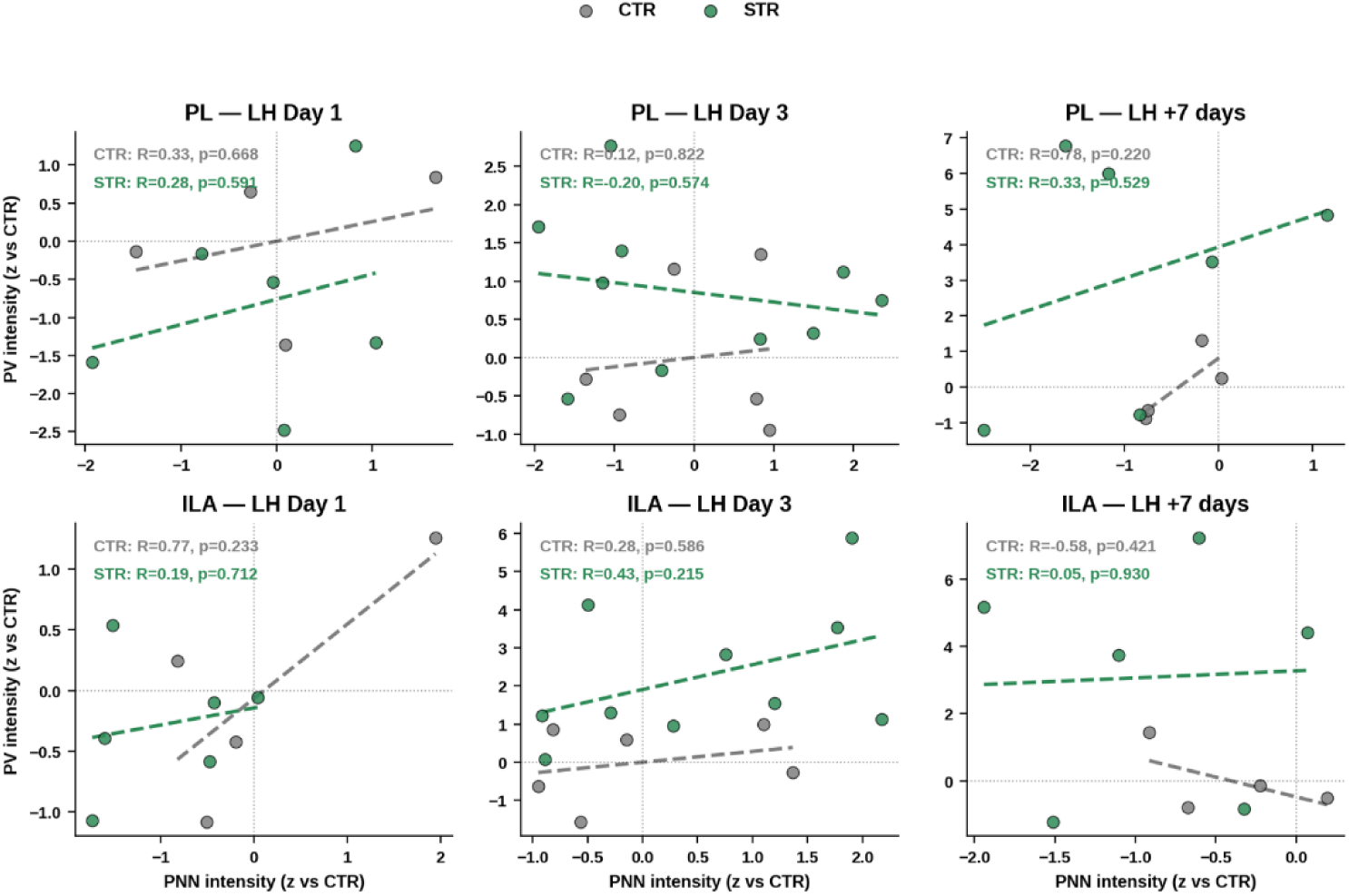
PV-PNN intensity couple across cohorts. Correlation between PNN and PV intensity per cohort: LH Day1, LH Day 3 and LH +7 days, in PL and ILA. Points represent individual animals from CTR (grey) or STR (green) animals, and dashed lines show group-specific linear fits. Correlations were evaluated using two-sided Pearson tests.

